# Rewired gene interactions during development of serially homologous appendages in male and female *Drosophila*

**DOI:** 10.1101/2024.10.19.618997

**Authors:** Amber M. Ridgway, Javier Figueras Jimenez, Beñat Yáñez Iturbe-Ormaeche, Maria D. S. Nunes, Alistair P. McGregor

## Abstract

*Drosophila* ventral appendages are considered to be serially homologous derived from a ventral appendage ‘ground state’ and shaped by different Hox inputs. In the legs and antennae a combination of the transcription factors C15, LIM1 homeobox 1 (Lim1), and Al (Aristaless) is required for the development of the tarsal claws and aristae, respectively. However, the roles of these factors in genital development have remained unexplored. Here, we investigated the expression and function of C15, Lim1, and Al in the development of male and female terminalia (genitalia and analia). We found that C15 plays distinct roles in males and females, repressing male clasper bristle formation while promoting bristle development in the female epiproct. Unlike in the antennal and leg discs, C15, Lim1, and Al are not all simultaneously co-expressed in any anal or genital structures in either sex, indicating that the interactions among these factors have diverged across these appendages. Nevertheless, we inferred regulatory interactions between C15 and other factors, reflecting similarities between leg and male clasper development. Finally, we identified a male-specific *C15* enhancer that is active in male claspers but not in the female epiproct, legs or antennae. This *C15* enhancer modularity may underpin tissue- and sex-specific regulatory logic.

## Background

The diversification of serially homologous structures has been a major theme during animal evolution (1, 2). This is exemplified by the elaboration of ventral appendages in arthropods that have evolved into a ‘toolkit’ of functionally distinct structures, from walking legs and swimming appendages to antennae, respiratory organs, spinnerets, and genitalia (1-6). It is thought that appendage diversity is regulated by distinct Hox gene inputs as well as other differences in the underlying developmental gene regulatory networks (GRNs) (4, 5, 7, 8). Understanding the regulation and diversification of these mechanisms is a key question in evolutionary developmental biology.

In *Drosophila,* the legs are patterned by overlapping expression domains of the transcription factors *homothorax* (*hth)*, *dachshund* (*dac*), and *Distal-less* (*Dll*), along the proximo-distal axis, with expression of the latter two genes overlapping in the distal tibia and tarsal segment (9) (Fig. 1). The distal part of the leg includes the tarsus, which is composed of five tarsal segments followed by a pre-tarsal region bearing a pair of claws (Fig. 1). Development of the tarsus and pre-tarsal region is also regulated by a proximo-distal gradient of increasing Epidermal growth factor receptor (EGFR) activity, which results in the activation of specific transcription factor-encoding genes, including *Bar* (*B*) and *bric a brac 2* (*bab2*), in the distal tarsal segments and *C15* (also called *clawless*) in the pretarsal region (0, 11). C15, in turn, activates *aristaless* (*al*) and *LIM homeobox 1* (*Lim1*), which feeds back to promote *C15* expression in the pre-tarsal region (0, 11). Together, C15, Lim1, and Al combine to repress *B* and *bab2* in the pre-tarsal region, with C15 and Al cooperatively binding to directly repress *B* expression and C15 competing with the activator Dll to regulate *bab2* expression (0-13). Loss of C15, Lim1, or Al expression in the leg results in the loss of the pre-tarsal region, including the claws (0, 11). This trio of transcription factors is also co-expressed and required for the development of part of another ventral appendage, the aristae of the antennae (0, 11, 14, 15). However, in developing eyes, only C15 and Lim1 are expressed in the ocellar region (0, 11, 14, 15). Therefore, although the combinatorial expression of C15, Lim1, and Al can differ according to the developmental context, these three transcription factors are essential for appendage patterning.

**Figure 1.**
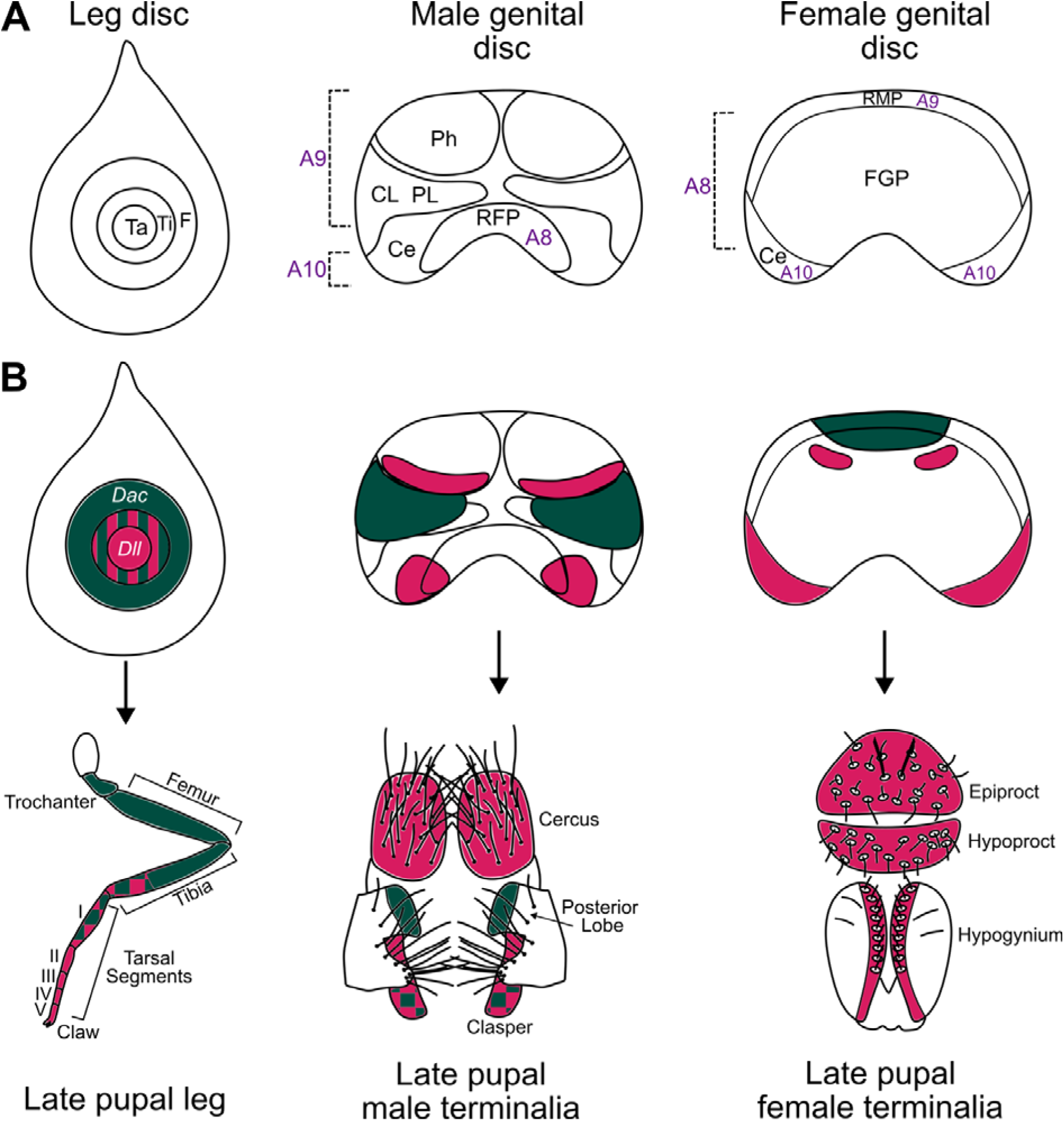
Development of the *Drosophila* leg and male and female terminalia. (A) Fate maps of the leg, male genital, and female genital discs. Ventral view of genital discs. The three segmental primordia of the terminalia (A8, A9 and A10) are indicated. (B) In the leg disc, *Dll* (pink) is expressed at the centre, which gives rise to the tarsal segments and distal tibia. *da*c (green) expression marks the first tarsal segment, tibia, femur, and trochanter. In the male genital disc, *Dll* is expressed in the phallic and cercus primordia. *dac* expression in the male genital disc marks the clasper and posterior lobe primordia. During male pupa development, *Dll* is expressed throughout the claspers and cerci, and *dac* is expressed in the proximal clasper region and posterior lobe. In the female genital disc, *Dll* is expressed in the cerci primordia and part of the genital primordium. *dac* is expressed in the repressed male primordium of the female genital disc. During female pupal development, *Dll* is expressed in both anal plates and part of the external female genitalia. *dac* is not expressed in the external structures, but instead localizes to internal genital structures (not shown). Note, that the internal phallic structures are not shown in B for simplicity. Diagrams are not to scale. Ta: Tarsus, Ti: Tibia, F: Femur, CL: Clasper, Ph: Phallus, PL: Posterior lobe, Ce: Cercus, RFP: Repressed female primordium, FGP: Female genital primordium, RMP: Repressed male primordium. The figure summarises previously published data (9, 17, 31, 35, 36).

Classic experiments in *Drosophila* suggested that the ‘ground state’ state of ventral appendages constitutes a leg-like proximal segment and distal tarsus (6). It was reasoned that inputs from Hox genes and *hth* pattern this basic structure to generate specific appendages, for example *Antennapedia* in the legs (6). In the genitalia, it is thought that the posterior Hox gene *Abdominal-B* (*Abd-B*) represses leg fate and promotes genital fate (7). Indeed, loss of *Abd-B* results in the development of tarsal-like structures from the genitalia in *Drosophila* and other insects, further supporting the ventral appendage ‘ground state’ model (6-18). While excellent progress has been made in understanding how Hox genes and downstream mechanisms regulate the identity and fine-scale morphology of appendages, especially legs, wings, and halteres in *Drosophila* (9-27), much remains to be learned about the development and evolution of genital structures.

The terminalia of *Drosophila* are composed of the analia and genitalia, which develop from the genital imaginal disc (8). This disc is composed of cells from abdominal segments A8, A9 and A10 (Fig. 1A). The analia of both sexes derive from segment A10 and give rise to the anal plates in males (cerci) and females (composed of the epiproct and hypoproct) (Fig. 1A) (8-30). In males, the genitalia, including the claspers (surstyli), epandrium and internal organs such as the phallus and associated structures, develop from A9 (8, 30) (Fig. 1A). In females, the genitalia develop from segment A8 (8, 29). Segments A8 and A9 form the repressed female primordium and repressed male primordium in males and females, respectively (8) (Fig. 1A). The primordia of adult terminal structures are marked by the expression of specific genes in the genital imaginal disc, for example *caudal* (*cad*) and *Abd-B* mark and regulate the development of the analia and genitalia, respectively (8). The regulation of analia and genitalia development also integrates sex-specific signals to specify and pattern male and female structures (8). During the late third larval instar (L3) and pupal stages, the cells of the anal and genital primordia divide, differentiate and undergo metamorphosis, causing these tissues to be remodelled and develop into the adult structures (8) (Fig. 1).

Comparisons of gene expression between genital and leg discs have revealed similarities consistent with the concept that the terminal structures are derived ventral appendages (7, 31, 32). For example, *Dll* expression in the four distal tarsal segments in the leg is reflected in the proximal domain of the clasper and throughout the cerci in the male genitalia, while overlap of *dac* and *Dll* in the first tarsal segment and part of the tibia, is similar to their expression in the distal clasper domain (Fig. 1B). Furthermore, it was shown that *C15* is expressed in the developing pupal claspers (3) and downstream target genes, including *bab2*, have also previously been shown to be expressed in the claspers (7, 33-35). However, the relative expression, functions, and interactions of key genes that regulate distal leg development in *Drosophila* have not previously been systematically studied in developing male and female terminalia. This information is crucial to not only understanding the development of these structures but also how they have been remodelled from the ventral appendage ‘ground state’ during evolution.

Here, we investigated the expression, function, and interactions of *C15* and upstream and downstream genes in the pre-tarsal GRN during the development of male and female *Drosophila* terminalia. Our results highlight differences in the deployment and interactions of these genes between the development of leg and terminalia, as well as between male and female terminalia development. Our data indicate that there has been extensive rewiring of the distal leg GRN during the evolution of male and female terminal structures, and although this re-programming involves Hox gene inputs, we present evidence that it has also potentially been underpinned by *C15* enhancer modularity.

## Materials and Methods

### Drosophila lines and manipulation of gene expression

*UAS-RNAi* lines were provided by the Vienna *Drosophila* Resource Centre (VDRC) (7) or the Bloomington *Drosophila* Stock Centre (BDSC) (NIH P40OD018537). All *UAS-RNAi* lines used are detailed in Supplementary Table 1. *UAS-RNAi* males were crossed to *NP6333-GAL4 (P{GawB}PenNP6333)* virgin females also carrying *UAS-Dicer-2 (P[UAS-Dcr-2.D]*) (4, 37, 38). All crosses were carried out at 25°C. Crosses were transferred to standard cornmeal food every two days and maintained under a 12-hour light/dark cycle. Progeny were aged at least 3 days and collected and stored in 70% Ethanol (EtOH) at -20°C.

The cerci, posterior lobes, and claspers of the adult male genitalia, and the epiproct, hypoproct, and hypogynium of adult female genitalia, were dissected using 0.14 mm diameter stainless steel pins in Hoyer’s solution (9). Dissected adult structures were mounted in Hoyer’s solution on slides containing eight individual 6 mm diameter wells. To account for body size, the T2 legs of each fly were also dissected and mounted. A Zeiss Axioplan light microscope with a Jenoptik ProgRes C3 camera was used to image each dissected structure, where 250X magnification was used for the terminal structures and 160X magnification for the T2 legs. The area of the posterior lobe, cercus, lateral plate, hypoproct, epiproct and length of T2 tibia leg were measured manually using ImageJ (0). When measuring the hypoproct and epiproct area, an outline was drawn and enclosed with an artificial baseline (1). Bristles on structures were counted under a light microscope using a tap counter. The area, bristle count, and length were recorded for both structures of each pair per individual, and the average was then used for the statistical analyses.

### Statistical analyses

All statistical analyses were carried out using R version 4.2.0 (2). Measurements of each structure described above were first assessed for normality using the Shapiro Wilk test. Normally distributed data were analysed using Dunnett’s test and ANOVA, whereas the Kruskal Wallis test was used for non-normally distributed data (3). RNAi knockdown progeny were compared to both parental line controls. If this test was statistically significant, the Tukey’s Test / Wilcoxon Rank Sum Test (BH p-adjusted method (4)) was used to identify whether the effect was significantly different to both parental controls (Supplementary Tables 1 and 2). The RNAi was considered to have had no effect if p =/> 0.05 or if p < 0.05 where the trait value in the RNAi knockdown progeny was intermediate to the two parental controls (Supplementary Tables 1 and 2). Where the T2 tibia length of the RNAi knockdown was significantly different to both parental controls following the tests described above, Pearson/Spearman correlation analysis was performed on the cercus, epiproct and hypoproct area dependent on Shapiro Wilk test results. If statistically significant, the area of the structure was divided by the tibia length squared, and statistical tests were carried out on the normalised measurements.

### Scanning electron microscopy

Scanning electron microscopy (SEM) was carried out on *NP6333-GAL4* and *NP6333-GAL4* > *UAS-C15* RNAi knockdown male and female terminalia. Flies initially stored in 70% EtOH were moved into 100% EtOH at least 24 hours prior to imaging. The posterior of the fly was dissected in 100% EtOH. Samples were processed in a critical point dryer and mounted on SEM stubs, then gold-coated for 30 seconds. The terminalia were imaged using SE mode at 5 kV in a Hitachi S-3400N SEM, with a working distance of 13 to 14 mm. The entire terminalia was imaged at a magnification of 250X, and individual genital and anal structures at 900X.

### Sample preparation and immunohistochemistry

3^rd^ instar larvae were collected in ice-cold 1 X PBS, cut in half, and inverted. Larval tissues were then fixed in 4% formaldehyde in 1 X PBS for 20 minutes at room temperature. Samples were rinsed in 1 X PBS with 0.1% TRITON-X-100 (PBT) and then washed in 0.3% PBT for 30 minutes. Following this, samples were blocked for 20 minutes in 10% normal goat serum (NGS) in 0.1% PBT. Larval samples were then incubated with the primary antibody overnight at 4°C (Supplementary Table 3). The next day, samples were washed for 5 minutes five times in 0.1% PBT. The secondary antibody was added and incubated at room temperature for 2 hours. Samples were washed accordingly, DAPI stained, and then dissected on slides to isolate the required imaginal discs. Discs were mounted in VECTASHIELD and sealed with a coverslip for imaging.

Male and female white prepupae were defined as 0 hours after puparium formation (hAPF). Males were identified by two circular transparent cuticles on either side of the posterior abdomen (5), and then stored at 25°C until the developmental timepoint required. Using micro-scissors, staged pupae were cut near the posterior end in ice-cold 1 X PBS, then flushed with 1 X PBS to remove as much fatty tissue as possible, followed by fixation in 4% formaldehyde for 30 minutes at room temperature. Samples were washed in 1 X PBS, then gradually brought to 100% EtOH for storage at -20°C. Before staining, the tissues were removed from the pupal casing, leaving just the exposed genital tissue samples enclosed in the outer membrane. All steps following this were carried out in 0.3% PBT and blocked and stained in 2% NGS in 0.3% PBT. Pupal samples were blocked for 2 hours, then incubated overnight at 4 °C with the primary antibody. The next day, pupal samples were washed for 6 x 20-minute intervals before the addition of the secondary antibody and then incubated at 4°C overnight. Samples were subsequently washed and DAPI stained if required, then the outer membrane was removed using dissection forceps prior to mounting. Microscope slides with coverslips used as walls were used to create a well to protect the sample from being crushed during imaging. Poly-L-Lysine solution was added to the well prior to mounting to avoid movement of the tissue. All pupal samples were mounted in Hydromount, and a coverslip was placed on top and sealed with nail varnish.

For all stains, the primary and secondary antibody concentrations are detailed in Supplementary Table 3. Where annotated as pre-absorbed primary antibody (Supplementary Table 3), the working concentration of the primary antibody was diluted in 0.3% PBT for pupae, and 0.1% Tween-20 in 1 X PBS for larval tissues, before being incubated with inverted, fixed larvae for 1 hour at room temperature, then stored at 4°C.

All stained samples were imaged using a Zeiss LSM800 upright Confocal Laser Scanning Microscope. A 20X air objective was used to image larval disc samples, and a 40X oil objective for pupal samples. Some pupal structures were imaged using a 63X oil objective when capturing individual external structures. Z-stacks with 0.7 μm and 1 μm separation were used for pupal and larval samples, respectively. All images were processed with Fiji (6).

To screen for active *C15* genital enhancers, males of available GAL4 enhancer reporter lines (stock numbers BL41233, BL50056 and BL50087) (7) encompassing intronic regions of the *C15* locus were crossed to *UAS-GFP* virgin females. To assess activity in live pupae, the outer cuticle was removed using dissection forceps, and pupae were mounted on a slide in a drop of 80% glycerol for imaging. Dorsal and ventral images were captured using a Zeiss Axiozoom microscope with a GFP filter and Z stacks of 1 μm separation.

### ATAC-seq library preparation

Prepupae were collected from *D. melanogaster* (Oregon R) and aged until 30 hAPF. Pupae were dissected in ice-cold 1 X PBS, followed by incubation in lysis buffer (10 mM Tris-HCL, pH 7.5; 10 mM NaCl; 3 mM MgCl_2_; 0.1% IGEPAL and 0.1% digitonin). After the lysis was stopped, the nuclei were collected by centrifuging at 500 g for 10 minutes. 50,000 nuclei were resuspended in Tagmentation Mix (25 µl tagmentation buffer (20 mM Tris-CH2COO-, pH 7.6; 10 mM MgCl_2_; 20% Dimethylformamide); 2.5 Tn5 transposase; 22.5 µl water) and incubated for 30 minutes at 37°C. 3 µl of 2 M NaAC, pH = 5.2 was added to the sample and then purified using a QIAGEN MinElute Kit. Library amplification was performed using NEBNext High Fidelity 2X PCR Mix with primers indexed with primers compatible with Nextera adaptors from Diagenode. 150 bp paired-end sequencing was carried out by Novogene (UK).

### ATAC-seq library, peak calling and footprinting

Raw reads generated from 30 hAPF male pupae were mapped to *D. melanogaster* genome version r6.55 Dec 2023 (8) using Bowtie2 (version 2.4.5) (9). Samtools (version 1.9) (0) was used to convert sam files to bam files and remove duplicates. Bedtools (version 2.31.1) (1) was used to convert generated data files into bed format, and reads were re-centered as previously described (2). Peak calling was then performed using MACS2 (version 2.2.7.1) (3) with the parameters -extsize 50, -q 0.01. ATAC-seq datasets for *D. melanogaster* eye-antennal imaginal disc (120 hours after egg laying) and pupal T2 leg were obtained from (4) and (1), and analysed as above.

Transcription factor footprinting analysis was carried out using the package TOBIAS (version 0.16.1) (5). Reads were corrected using ATACorrect, which corrects the bias of Tn5 transposase for specific sequences, and this correction was then used to score regions within peaks for the likelihood of being bound by proteins. We searched for motifs within the footprints and compared them against the JASPAR core (6) and CIS-BP (7) databases to identify putative transcription factors that bind to these sites (Supplementary Tables 4 and 5).

### Data availability

ATAC-seq data accession numbers: GSE279524 (pupal genital data), GSE113240 (T2 pupal leg data), and PRJNA666524 (eye-antennal-disc data).

## Results

### C15 expression in developing male and female terminalia

It was previously shown that *C15* mRNA is expressed in the proximal region of male claspers during pupal stages (3, 58). We examined the clasper expression of C15 further during male larval and pupal genital development by staining with anti-C15 and co-staining with anti-Odd-paired (Opa), which is known to mark the developing claspers (3) (Fig. 2A-A’’). We found that C15 protein is localised to a subset of cells within the Opa-marked clasper primordium of male larval genital discs (Fig. 2A-A’’). During pupal stages, C15 remains localised to the clasper but is restricted to cells in the proximal region that do not develop bristles, consistent with *C15* mRNA expression (3) (Fig. 2B-D). In addition, C15 expression extends from the proximal domain of the claspers to the periphery of the cerci from as early as 26 hAPF (Fig. 2B-D). Throughout pupal development, C15 is also detected in the abdominal tissue dorsal to the male terminalia (Fig. 2B-D).

**Figure 2.**
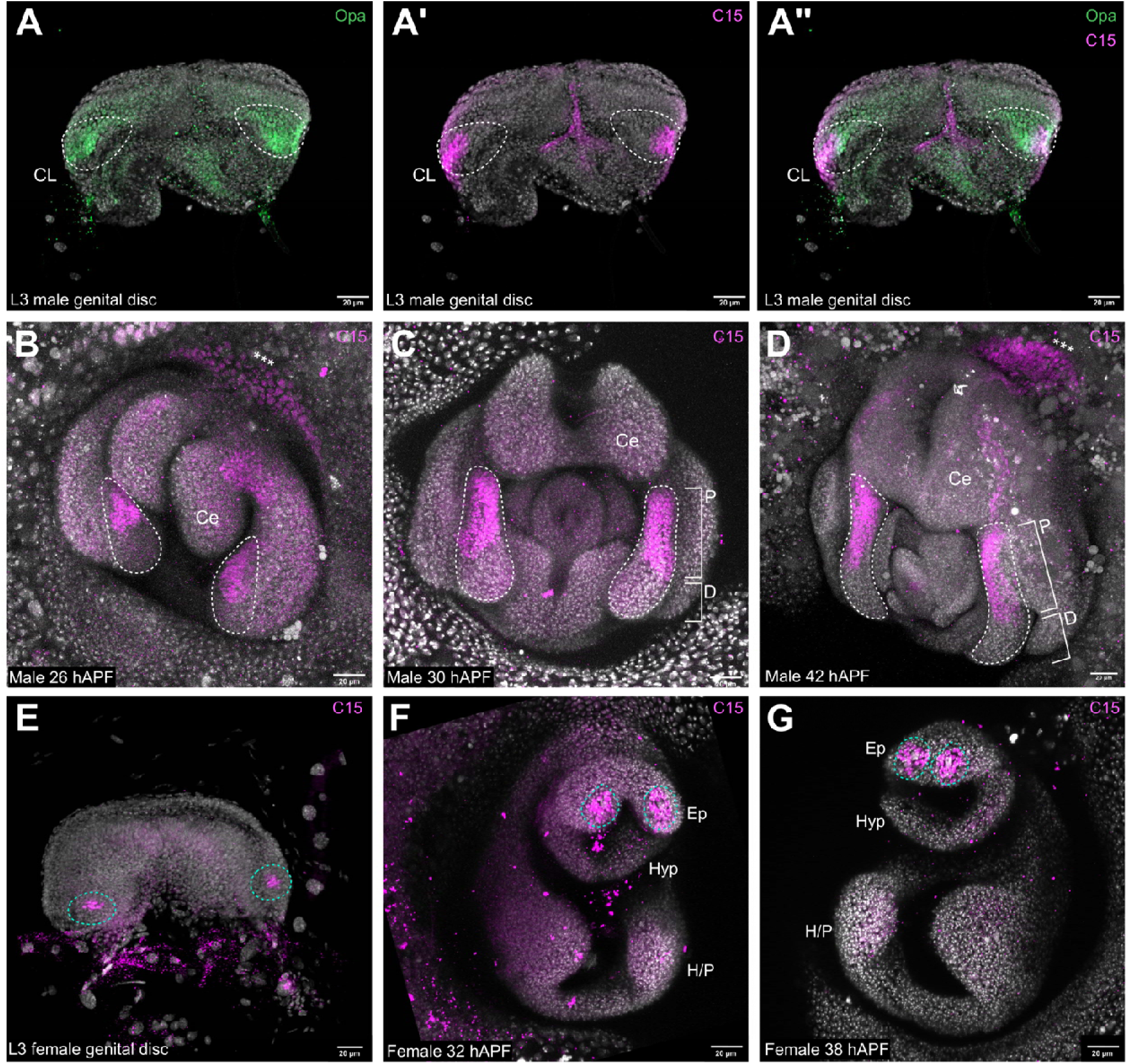
C15 expression in developing male and female terminalia. (A-A’’) During male genital disc development, C15 is expressed in the clasper primordium (CL), as confirmed by overlap with Opa. (B-D) During male terminalia pupal development, C15 is expressed in the proximal region of the clasper, peripheral to the cerci (Ce) and at the adjoining terminal tissue (marked by ***). (E) C15 is expressed in a specific cluster of cells within the female cerci primordia. (F,G) During female terminalia pupal development, C15 is expressed in the epiproct (Ep), encircling two prominent bristle progenitor cells. White dashed outlines mark the developing male claspers. Cyan dashed outlines mark C15 expression in the female epiproct. Proximal (P) and distal (D) domains of the male pupal claspers are labelled in C and D. All genital discs are the ventral view. Hyp: Hypoproct, H/P: hypogynium and oviprovector primordia. All samples were counterstained with DAPI.

We next investigated C15 localisation in the developing female terminalia. In females, C15 is expressed in two well-defined cell clusters of approximately seven cells in the genital disc (Fig. 2E). This expression pattern is maintained during pupal development and surrounds the two prominent sensilla of the epiproct consistent with *C15* mRNA expression (8) (Fig. 2F,G).

### C15 is required for bristle development in male and female developing terminalia

We then examined the function of *C15* during male and female terminalia development using RNAi knockdown. In males, we found that knockdown of *C15* expression resulted in a 17% increase in clasper bristle number (Fig. 3A; Supplementary Table 1). The extra ectopic bristles developed in the internal proximal domain of the clasper, a region that normally only displays bristles along the outer periphery (Fig. 3C-D’) (9), consistent with C15 expression in this region (Fig. 2C,D). The Hox gene, *Abd-B*, is expressed in all structures of the male genitalia and dictates tissue identity (Supplementary Fig. 1A). RNAi against *Abd-B* appeared to only produce a weak knockdown because the genitalia still developed relatively normally, but we did observe a significant decrease in clasper bristles (Supplementary Fig. 1B). Consistent with this, *Abd-B* RNAi knockdown appeared to extend C15 expression somewhat into the distal clasper domain, as well as generating stronger C15 expression surrounding the cerci (Supplementary Fig. 1C; Supplementary Table 1). Together, these results suggest that C15 represses clasper bristle development proximally and that this localisation of *C15* may be regulated in part by Abd-B repression.

**Figure 3.**
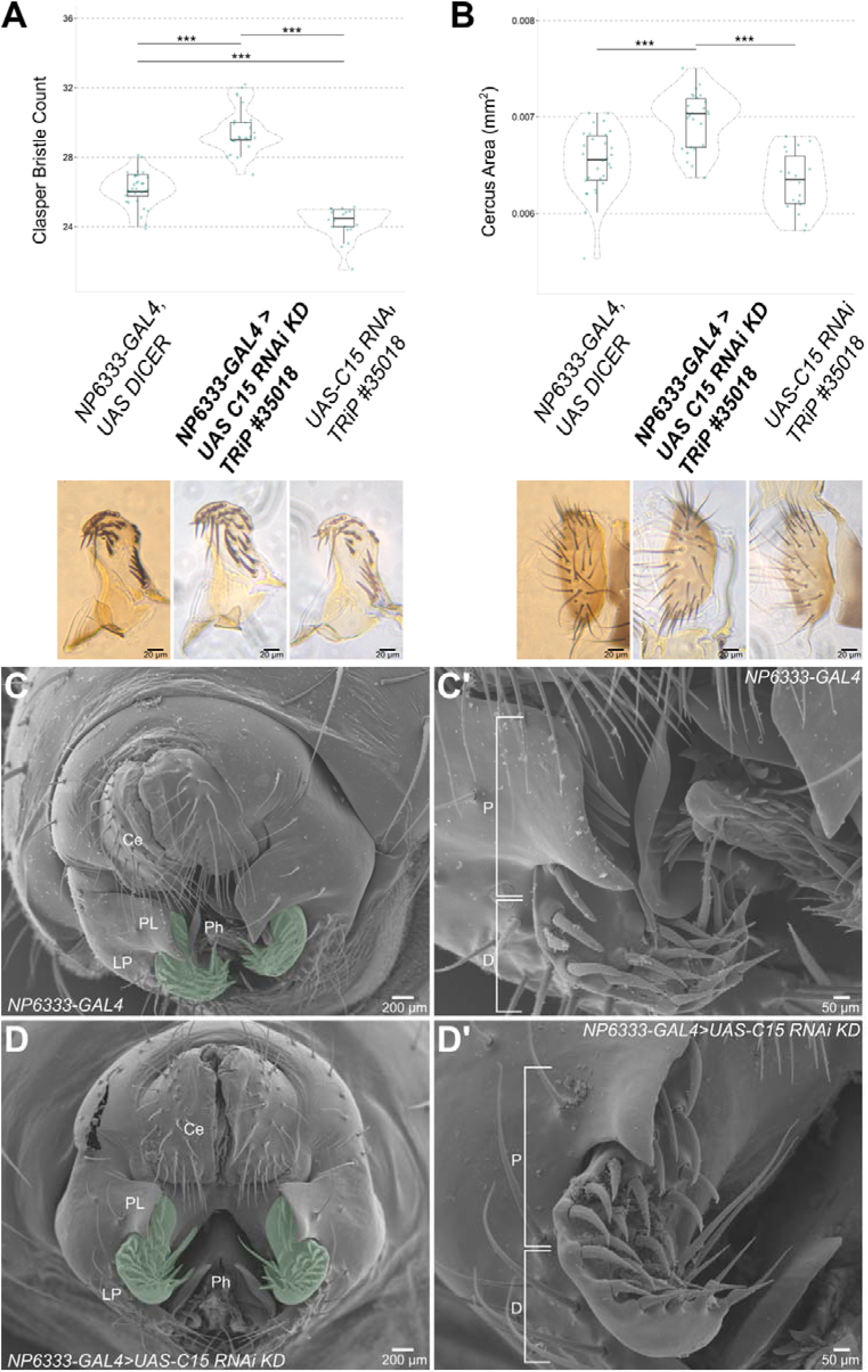
RNAi knockdown of *C15* during male terminalia development. (A) RNAi knockdown of *C15* in the developing male terminalia increased clasper bristle number, and (B) increased cercus size (Ce). SEM images of *NP6333-GAL4* parental control (C,C’) and of *NP6333-GAL4>UAS-C15* RNAi knockdown (D,D’). *** p < 0.001, (n > 17) (Supplementary Table 1). (C, D) Green shading marks the claspers. (C’,D’) Proximal (P) and distal (D) regions of the clasper are indicated. PL, posterior lobe. LP, lateral plate. Ph, phallus.

RNAi knockdown of *C15* also significantly increased the area of the male cerci (Fig. 3B) (Supplementary Table 1), although this gene is only expressed at the boundary of this structure (Fig. 2D). However, no difference in cerci bristle number was detected upon *C15* knockdown (Supplementary Fig. 2). Surprisingly, male T2 tibia length also increased upon *C15* knockdown even though this gene is not expressed in the leg disc corresponding to this part of the leg (Supplementary Fig. 2).

In females, *C15* RNAi knockdown eliminated the two prominent epiproctal sensilla (Fig. 4A,A’,C,C’), consistent with the expression of this transcription factor (Fig. 2F,G). However, *C15* RNAi knockdown in females had no significant effect on the total number of surrounding epiproct bristles compared to controls, although the area of the epiproct did significantly decrease (Fig. 4B,B’; Supplementary Table 2). *C15* RNAi knockdown had no detectable effect on the hypoproct or hypogynium bristle patterning, and unlike in males, there was no significant effect on female tibia length (Supplementary Fig. 2). In summary, these results revealed an epiproctal-specific role for *C15* in females, where it appears necessary for the development of the two prominent epiproctal sensilla and epiproctal size.

**Figure 4.**
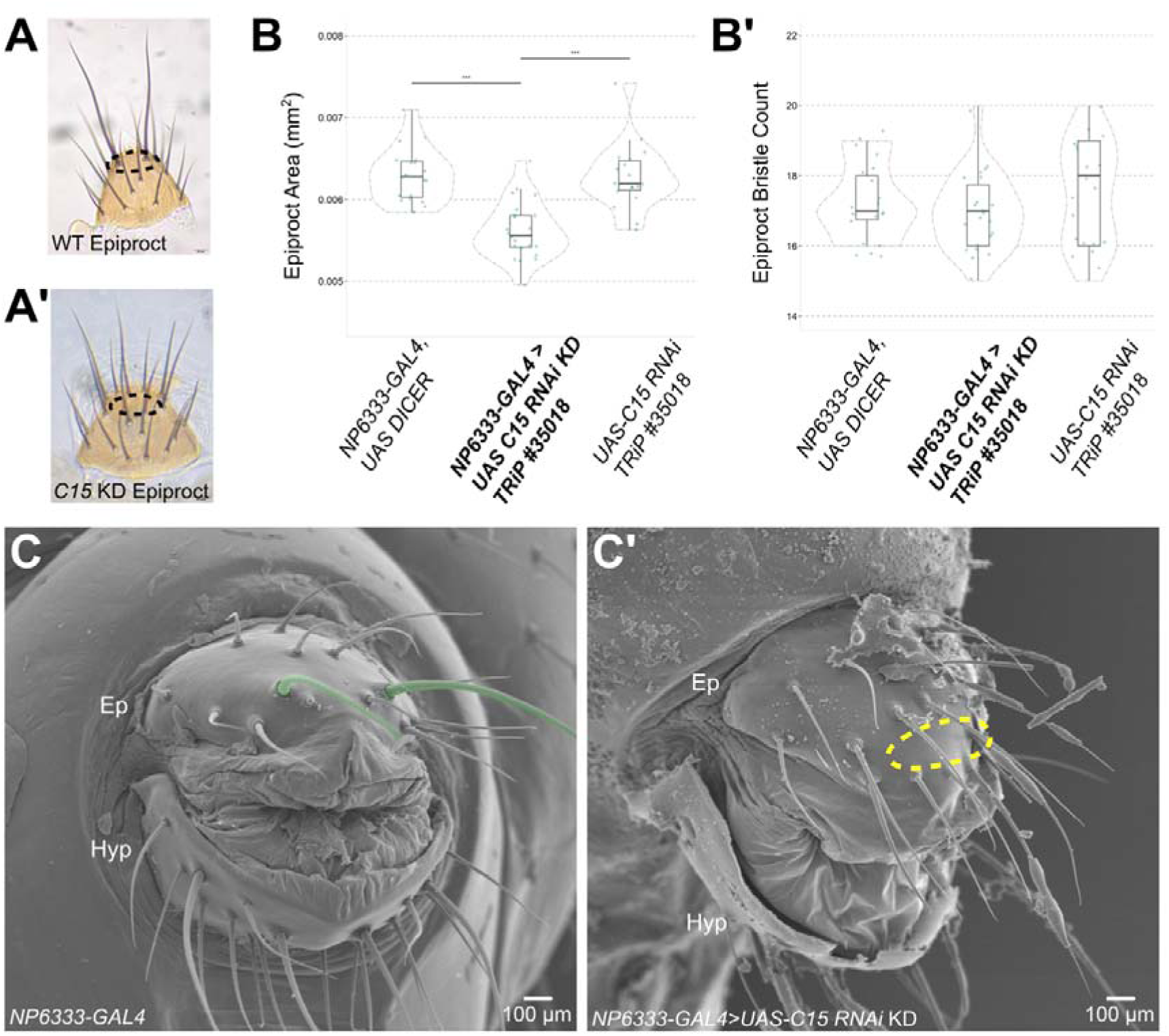
RNAi knockdown of *C15* during female terminalia development. (A,A’) Light microscopy images of the female epiproct, where the dashed black ovals mark the region of the two prominent epiprocta sensilla. (B) RNAi knockdown of *C15* significantly reduced the epiproct area, but did not affect the total epiproct bristle count (B’). *** p < 0.001, (n > 17) (Supplementary Table 2). (C) SEM image of *NP6333-GAL4* parental control. The two prominent epiproctal sensilla are coloured green. (C’) *NP6333-GAL4>UAS-C15* RNAi knockdown female terminalia. The dashed yellow oval marks the loss of the two longer epiproctal sensilla. Ep: Epiproct and Hyp: Hypoproct.

### Analysis of other leg transcription factors relative to C15 during terminalia development

We next investigated other key transcription factors in the terminalia that are part of the distal leg GRN involving *C15*. In the developing leg, *dac* is expressed proximally up to the first tarsal segment but is absent from the most distal tarsal segments and the pre-tarsal region (0) (Fig. 1B). It was previously shown that *dac* is also expressed in the clasper primordia of the male genital disc (Fig. 1B) (7). We used a hypomorphic allele of *dac* with a GAL4 insertion crossed to *UAS-GFP* to assay *dac* expression in pupal genitalia (1). We observed that *dac* expression predominantly localises to the distal clasper and lateral plates, with weaker expression in the proximal domain of these structures (Supplementary Fig. 3A). Consistent with this, the claspers of the *dac* hypomorphic mutants exhibited loss of clasper bristles and the remaining distal bristles appear elongated (Supplementary Fig. 3B; Supplementary Table 1). These findings suggest that *dac* promotes male clasper bristle development but may also regulate bristle morphology.

Dll is expressed throughout the distal leg segment primordia, but its direct activation of *bab2* is blocked by C15 in the pre-tarsal region (2). In the male genital disc and pupal terminalia, *Dll* is expressed in the cerci and phallic primordia (1) (Fig. 1B and Supplementary Fig. 3C). We found that Dll was also expressed in both the proximal and distal regions of the claspers during pupal development (Supplementary Fig. 3C). RNAi knockdown of *Dll* resulted in a small increase in clasper bristle number (Supplementary Fig. 3B), and reduced cerci size and bristle count, consistent with previous work (Supplementary Table 1) (1). However, we did not detect any obvious effect of *Dll* RNAi knockdown on the expression of C15 (Supplementary Fig. 3D). Interestingly, like in the leg disc, *bab2* and C15 expression do not overlap in the male genital disc (Supplementary Fig. 4A,B), nor in pupal genitalia where *bab2* is only expressed in the distal clasper (and in the cerci) (Supplementary Fig. 4). This suggests C15 may inhibit Dll-mediated activation of *bab2* in the proximal clasper like in the leg pre-tarsal region (2).

In leg imaginal discs, *C15* expression overlaps with *Lim1* and *al* in the leg pre-tarsal primordia, and throughout pupal leg development (Supplementary Fig. 5) (0, 11). In this context, these three transcription factors co-regulate one another, which is essential for tarsal claw development (0, 11, 15). For example, loss of *C15* reduces *al* expression and vice versa, and loss of either *al* or *C15* abolishes *Lim1* expression, while *C15* expression is markedly reduced in *Lim1* mutants (0, 11, 15). We therefore assayed their relative expression during terminalia development. We did not detect any Al expression in the developing male claspers during larval or pupal stages (Fig. 5A,A’). Consistent with this, *al* RNAi knockdown showed no detectable clasper phenotype and did not affect C15 expression (Supplementary Fig. 6).

**Figure 5.**
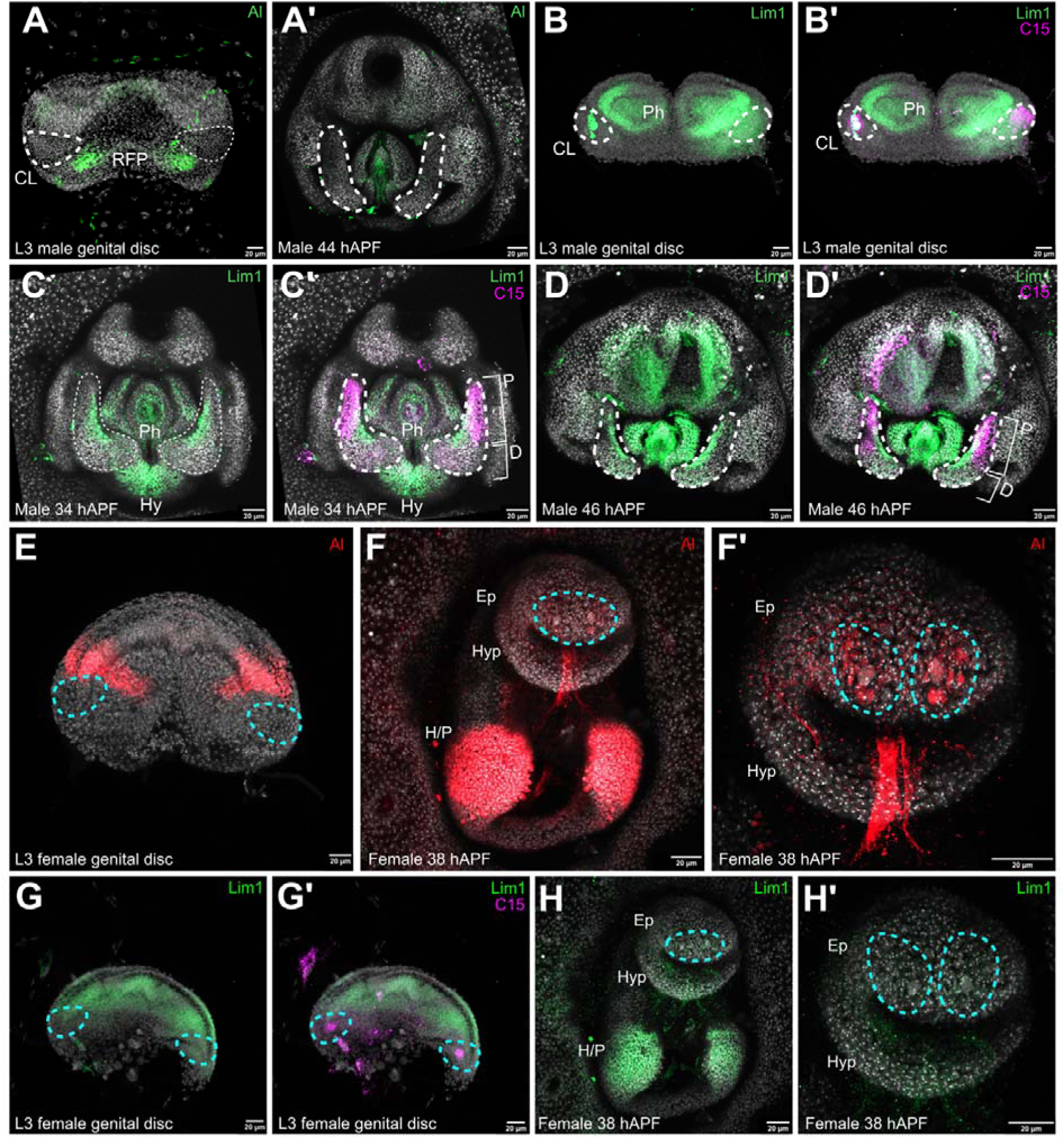
Expression of Al and Lim1 in the male and female developing terminalia. Al is expressed in the repressed female primordium (RFP) of the male genital disc but not in the primordia of any genital structures in the disc or during pupal stages (A,A’). (B,B’) Lim1 is expressed in the phallic region (Ph) of the male genital disc, and partially overlaps with C15 expression in the clasper primordia (CL). (C-D’) During male pupal development, Lim1 is expressed in the inner region of the proximal (P) and distal (D) clasper domains, where it partially overlaps with C15 proximally. Lim1 is also expressed throughout the phallic structures and hypandrium (Hy). (E) In female genital discs, Al is expressed in a defined region of the hypogynium/oviprovector (4). (F) During female pupal development, Al is expressed in the female hypogynium and oviprovector primordia (H/P) and epiproct (Ep) ventral to the two prominent epiproctal sensilla (F’). (G,G’) Lim1 is expressed in the hypogynium/oviprovector primordia of the female genital disc, but does not overlap with C15 expression in the cerci primordia. (H,H’) Lim1 is only expressed in the hypogynium and oviprovector primordia (H/P) of female pupal terminalia. Hyp: Hypoproct. (A-D’) Dashed white outlines mark the clasper primordia. (E,H’) Dashed cyan ovals mark the epiproctal C15 expression region. All samples were counterstained with DAPI.

In the female genital disc, Al is expressed in the hypogynium/oviprovector primordia but not in the epiproctal primordium marked by C15 (Fig. 5E). Post-metamorphosis, Al expression continues in the hypogynium/oviprovector and is also detected in the epiproct ventral to the two prominent sensilla (Fig. 5F,F’). Note that while it appears that Al and C15 are expressed in the same region of the epiproct, we were unable to determine the extent of their overlap because both antibodies originated from rat. *al* RNAi knockdown reduced epiproct bristle count and area, although the two prominent sensilla specified by C15 were still present (Fig. 6A). Knockdown of *al* expression resulted in loss of C15 in the cells ventral to the epiproctal sensilla (Fig. 6C-D’), where they overlap in expression (Fig. 5). *C15* RNAi also completely abolished Al expression in the epiproct (Fig. 6E-F’). Taken together these results confirm that there is positive regulation between *al* and *C15* in the female epiproct like that seen in the leg pre-tarsal region (0, 11, 13).

**Figure 6.**
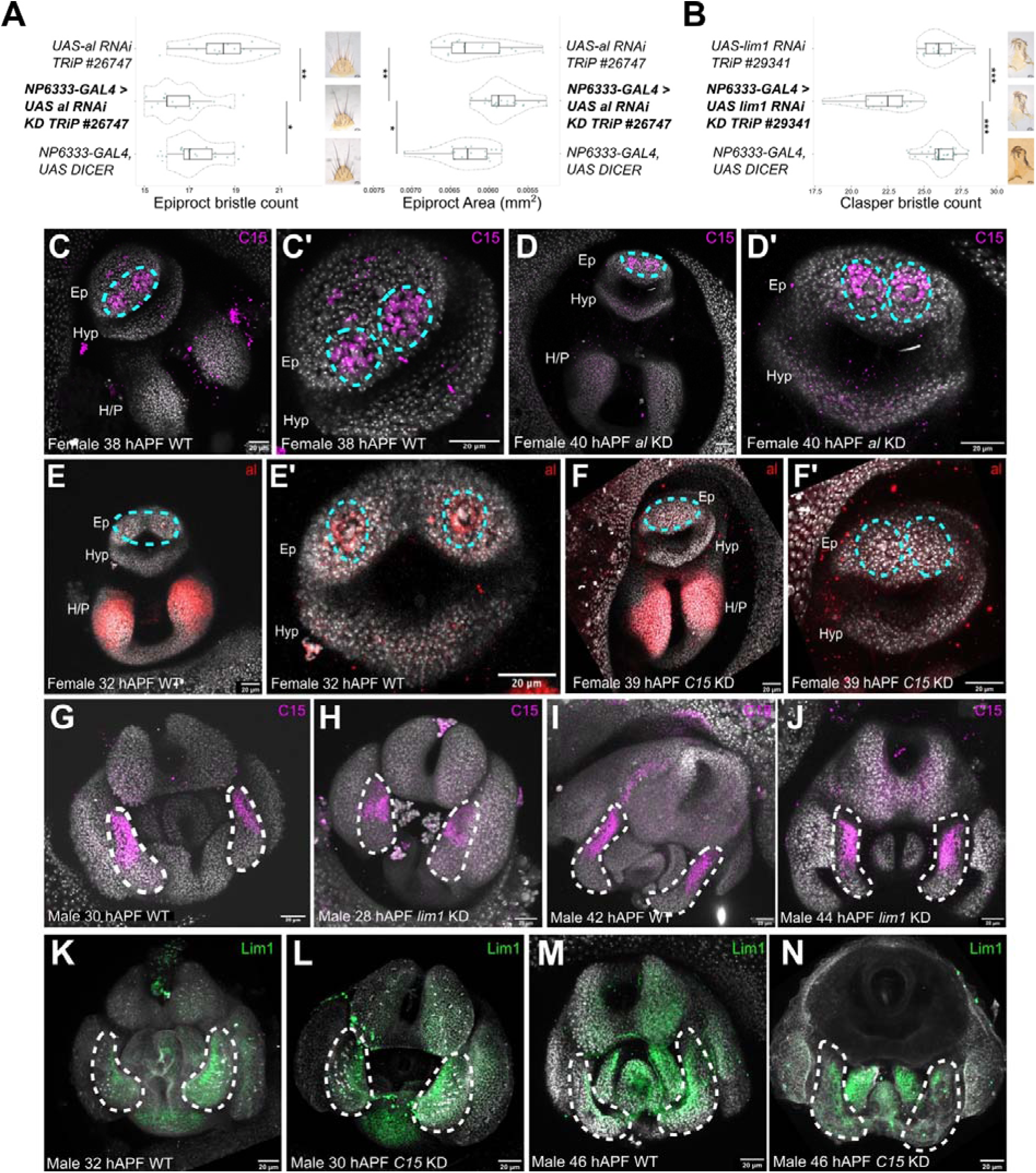
Roles of *Lim1* and *al* during male and female terminalia development. (A) In females, *al* RNAi knockdown reduced epiproct bristle count, and area. (B) In males, *Lim1* RNAi knockdown significantly reduced clasper bristle count. *** p < 0.001, ** p < 0.01, * p < 0.05 (Supplementary Tables 1 and 2). n > 14 for all lines phenotyped. (C,C’) C15 expression in the female pupal epiproct. (D,D’) *al* RNAi knockdown abolished C15 expression only in cells ventral to the female epiproctal sensilla primordia. (E,E’) C15 expression at the female pupal ovipositor and epiproct. (F,F’) *C15* RNAi knockdown entirely abolished Al expression in the female epiproct. (G,I) C15 expression in the proximal clasper domain in developing male terminalia. (H,J) *Lim1* RNAi knockdown did not appear to affect C15 expression in male pupal terminalia. (K,M) Lim1 expression in the inner region of the clasper, spanning both the proximal and distal regions. (L,N) *C15* RNAi knockdown did not appear to affect Lim1 expression in male pupal terminalia. Ep: Epiproct, Hyp: Hypoproct, H/P: hypogynium/oviprovector. (C-F’) Dashed cyan ovals mark the region of epiproctal C15 expression. (G-N) Dashed white outlines mark the clasper primordia. All samples were stained with DAPI.

We observed Lim1 expression in male genital discs partially overlapping with C15 (Fig. 5G,G’). During pupal stages, Lim1 expression was maintained in the clasper and was also observed in the hypandrium and phallus (Fig. 5C-D’). At 34 hAPF, as the clasper and lateral plate primordia are separating, Lim1 is expressed in a thin strip of cells at the inner region of the clasper, partially overlapping with C15 in the proximal region and reflecting their earlier relative expression in the genital disc (Fig. 5C,C’). In older pupae, Lim1 clasper expression extends further distally retaining some overlap with C15 in the proximal domain (Fig. 5D,D’).

*Lim1* RNAi knockdown resulted in significantly fewer clasper bristles (Fig. 6B), opposite to the effect of *C15* RNAi (Fig. 3). Furthermore, unlike in the developing pre-tarsal region, there does not appear to be an obvious regulatory interaction between C15 and Lim1 in male genitalia because although their expression overlaps in a few cells in the developing claspers, RNAi against one had no detectable effect on the other’s expression pattern (Fig. 6G-N).

In developing female terminalia, there was no detectable Lim1 in the epiproct during larval and pupal development although, Lim1 was expressed in the hypogynium/oviprovector primordia like Al (Fig. 5H). As expected, *Lim1* RNAi knockdown in females did not affect epiproct bristle or area (Supplementary Fig. 6B). However, *Lim1* RNAi knockdown did decrease hypogynium bristle number, consistent with its expression pattern (Supplementary Fig. 6C,C’).

Overall, analysis of the leg pre-tarsal core trio of transcription factors in terminalia showed that in males, only Lim1 and C15 regulate clasper development, whereas in females, only C15 and Al regulate epiproct bristle patterning, while Lim1 and likely Al regulate hypogynium bristles without C15. These results contrast with the leg and aristae, which both employ all three of these transcription factors.

### A C15 genital enhancer is not active in the developing leg or arista

We next explored the cis-regulatory sequences that drive *C15* expression in the genitalia. Three GAL4 reporter lines spanning the first and second *C15* introns were available for this region (Fig. 7A) (7). We then generated ATAC-seq data for male pupal genitalia to compare the profiles of the *C15* locus, and in particular the regions captured in reporter lines, with T2 pupal legs and L3 eye-antennal discs (Fig. 7A). The intron 1 region spanned relatively inaccessible chromatin in the genital, leg, and eye profiles (Fig. 7A). The Intron 2 enhancer 1 region contained two peaks in the genital track that were also accessible in the other two tissues (Fig. 7A). The intron 2 enhancer 2 region, which overlaps with intron 2 enhancer 1 by 1007 bp, and encompasses the previously mapped embryonic enhancer of *C15* (2) (Fig. 7A), is relatively inaccessible in all three tissues.

**Figure 7.**
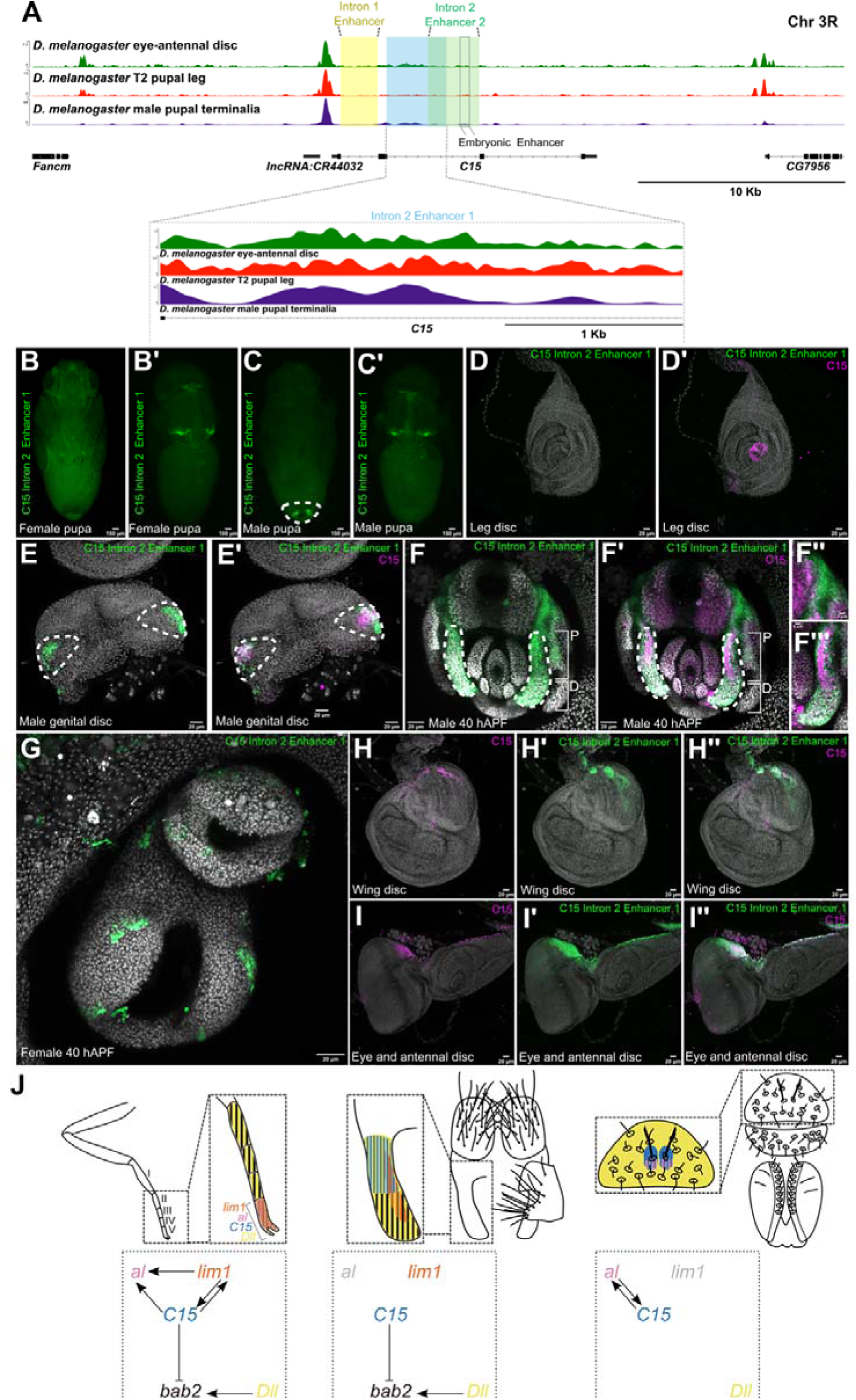
Activity of a *C15* male genital enhancer and regulatory interactions of this transcription factor in different developmental contexts. (A) *C15* locus ATAC-seq profiles in T2 pupal legs, eye-antennal discs, and male pupal terminalia. Regions encompassed by genitalia enhancer reporter lines are labelled accordingly. (B-C’) *C15* intron 2 enhancer 1 driving *UAS-GFP* expression in female (B,B’) and male (C,C’) late pupae. (B) In female pupae, C15 intron 2 enhancer 1 is not active in the ventral (B’) but drives expression in the wing hinges and near the ocelli in the dorsal. (C) *C15* intron 2 enhancer 1 is active in the male pupal terminalia (encircled by a dashed oval), (C’), and in the wing hinges and near the ocelli in the dorsal view. (D,D’) *C15* intron 2 enhancer 1 is not active in the leg disc. (E,E’) *C15* intron 2 enhancer 1 is active in male genital discs and overlaps with endogenous C15 expression in the clasper primordia. (F-F’’’) This enhancer is active in the claspers and periphery of the cerci in male pupal terminalia and overlaps with C15 expression in the proximal region of the clasper. (G) *C15* intron 2 enhancer 1 does not drive any specific expression resembling endogenous C15 expression in developing female terminalia. (H-H’’) This enhancer is active in the notum of the wing disc and overlaps with C15 expression. (I-I’’) *C15* intron 2 enhancer 1 is active in the cuticle and ocelli primordium of eye-antennal discs and overlaps with C15 expression. All samples have been stained with DAPI. P: proximal and D: distal clasper regions. J. Development of the pre-tarsal and tarsal segments is coordinated by a *C15*-centered GRN. *Lim1*, *al*, *bab2*, *Dll* and *C15* are expressed at the distal region of the leg where the claw protrudes from. *C15* and *bab2* expression are mutually exclusive to tarsal segments II-IV and the pretarsal region, respectively, while *Dll* spans the entire tarsus. *C15*, *Lim1* and *al* are all expressed in the pre-tarsal region and wired into an interdependent positive feedback loop. The entire male claspers and female epiproct are marked by *Dll* expression, like the tarsal segments of the leg. Similar to the leg, *C15* and *bab2* are expressed in mutually exclusive regions of the proximal and distal clasper, respectively. *Lim1* expression spans a thin inner strip of cells bridging the proximal and distal regions of the clasper. *al* is not expressed in the clasper. The female epiproct lacks *Lim1* expression. However, like the pretarsal region of the leg, *al* and *C15* show overlapping expression patterns. *bab2* expression in the female epiproct was not assayed in this study. Data summarised from this work and previous studies (9, 17, 31, 35, 36).

The *C15* intron 1 region did not drive expression in genital pupal structures or eye-antennal discs (Fig. 7A; Supplementary Fig. 7A,A’, F). However, this region was active in 3 or 4 leg disc cells some of which also expressed C15 (Supplementary Fig. 7C,C’). Female genital discs had no observable intron 1 enhancer activity, whereas the male genital disc did show reporter expression but in the repressed female primordium, which does not recapitulate endogenous *C15* expression (Supplementary Fig. 7D-E’).

Intron 2 enhancer 1 was active in the male terminalia but did not drive any specific expression in the female counterpart (Fig. 7). We detected intron 2 enhancer 1 activity in the clasper primordia of male genital discs, which was sustained during late pupal development (Fig. 7E-F’’’). This enhancer’s activity recapitulated the endogenous expression of *C15* in the proximal clasper, but was also active in the distal clasper where *C15* is not expressed (Fig. 7F’’’). Intron 2 enhancer 1 was not active in leg discs (Fig. 7D,D’) nor the arista primordia of the antennal disc, but was able to drive some expression in the ocellar primordium of the eye disc and notum in the wing disc (Fig. 7H-I”).

Intron 2 enhancer 2 lacked activity in genital, leg, and eye-antennal discs as well as the corresponding pupal tissues (Supplementary Fig. 7). However, this enhancer was active in the notum of the wing disc, which overlapped with endogenous C15 expression (Supplementary Fig. 7I) (3).

In summary, *C15* expression in the male genitalia is regulated, in part, by the intron 2 enhancer 1 region. Although this region also recapitulates aspects of the *C15* in the eye and wing discs, it does not drive expression in either the leg pre-tarsal region or arista primordia where *C15* overlaps with both *Lim1* and *al*.

We next used ATAC-seq footprinting to predict transcription factor binding sites in the intron 2 enhancer 1 region in the developing genitalia (Supplementary Tables 4 and 5). This analysis predicted high probability binding sites for Abd-B as well as other transcription factors expressed in the genitalia and analia, such as Daughterless and Cad (3), which could be candidates for activation or repression of *C15* expression in these developing tissues, respectively

(Supplementary Tables 4 and 5). Curiously, high probability binding sites were also detected for Al, but these may represent motifs bound by other homeobox-containing transcription factors because as our findings show that Al is not expressed in developing male terminalia (Supplementary Tables 4 and 5).

## Discussion

### C15 regulates bristle patterning on the female epiproct and male claspers

It was shown previously that C15 is vital for the development of the pre-tarsal claws, as well as the arista, ocellar postvertical bristles, and medial thorax bristles (0, 11). *C15* is also expressed in the developing male genitalia and female analia (3, 58) (this work), and we have shown that this transcription factor regulates bristle patterning on the male claspers and female epiproct. In females, C15 is required for the development of the two distinctive long epiproct sensilla consistent with the role of this transcription factor in promoting bristle development on other structures (0, 11). In contrast to promoting the development of female epiproct bristles, *C15* RNAi knockdown increased the number of male clasper bristles, indicating that C15 acts to repress bristle formation on these genital organs. These results suggest that C15 can play different roles in different developmental contexts that likely depend on which other transcription factors and co-factors it is expressed with in local GRNs.

### C15 GRN differences between male and female terminalia and among serially homologous appendages

C15, Lim1 and Al overlap in expression in the aristae of the antennae and pre-tarsal region of the legs and are crucial for the correct development of these structures (0, 11, 14, 15). However, we did not identify any male or female anal or genital structures where the expression of all three of these transcription factors overlapped. For example, we did not detect any Al expression in developing male claspers and only observed a small overlap in C15 and Lim1 expression in this developing organ (Fig. 7J). Furthermore, our results suggest there is no regulatory interaction between Lim1 and C15 in the developing male claspers, and that they appear to play opposing roles in promoting and suppressing clasper bristle development, respectively.

In the developing female genitalia, in the hypogynium/oviprovector primordia, we found an overlap between Lim1 and Al expression but no C15 expression. We did not observe expression of any of these transcription factors in the male analia, but C15 is required for Al expression in the female epiproct, although Lim1 is not expressed in this tissue (Fig. 7J). Therefore, the expression, roles, and interactions of C15, Lim1, and Al differ between male and female terminal structures. These results are consistent with other tissues that employ these factors individually or in pairs (0, 11, 14, 15, 62). Indeed, even in the pre-tarsus, for example, while Al and C15 combine to repress *Bar,* C15 is able to inhibit activation of *bab2* by Dll independently of Al, showing that even though their expression overlaps, they can have different roles (0-13).

Our results show that although the legs and genitalia are thought to be serial homologues, the deployment of C15-Al-Lim1 has diverged between these developmental contexts. Nevertheless, we did detect similarities in C15 expression and potential regulatory interactions with other factors between the claspers and the pre-tarsal region consistent with likely ancient serial homology. In the developing leg, *dac* is not expressed in the distal tarsal segments nor the pre-tarsal region where C15 is expressed, while *Dll* is expressed throughout the distal leg (9) (Fig. 7J). In the clasper, *dac* is only expressed in the distal region where there is no C15 expression while *Dll* is expressed throughout the developing clasper (Fig. 8). Intriguingly, *bab2* is expressed in the distal clasper which suggests that Dll activates this gene in this domain but *bab2* expression is blocked by C15 in the proximal clasper perhaps by inhibition of Dll activation like in the pre-tarsal region of the leg (2) (Fig. 7J).

Taken together, the relative expression and functions of *C15* and other genes suggest that *C15* patterns the proximal region of the clasper in a similar way to its role in the pre-tarsal region, but that there are key differences in wiring of the respective GRNs (Fig. 7J).

### Regulation of C15 expression in developing terminalia

We identified an enhancer that recapitulates *C15* expression in the developing male genitalia, although the activity of this cis-regulatory element extends into the distal region of the pupal clasper, where this transcription factor is not expressed. This ectopic activity somewhat resembles the expansion of C15 expression we observed upon *Abd-B* RNAi, which suggests that although we detected binding sites for this Hox factor in the *C15* enhancer, additional Abd-B sites may be needed to repress *C15* in the distal clasper. Other possible candidate transcription factors that may directly activate this enhancer, inferred from ATAC-seq footprinting, include Da and Cad, which could activate and repress *C15* in the male genitalia and analia, respectively (3). However, some of the most highly significant binding sites identified by JASPER and CIS-BP were for predicted homeobox binding targets, and given that these transcription factors can bind very similar motifs, further work is needed to identify which transcription factors directly regulate *C15* expression (Supplementary Tables 4 and 5). The *C15* genital enhancer we have identified also drives expression consistent with endogenous *C15* expression in the ocelli primordia of the eye-antennal disc and notum of the wing disc, but again, additional analysis is needed to identify the transcription factor binding sites responsible for the regulation of this element in these tissues and whether some of the same motifs are used in the genitalia or not.

Our results suggest that *C15* employs a different enhancer in developing female terminalia since the enhancer we identified does not drive expression of a *C15* pattern in the epiproct. It is also clear from our results that the previously characterised 650 bp embryonic *C15* enhancer (within the intron 2 enhancer 2 region), which is directly regulated by Zen and Smad (2), does not drive expression in either male or female terminalia, although this region is active in the wing notum (Supplementary Fig. 7I).

None of the putative *C15* enhancer fragments we tested, including the identified genital enhancer, fully recapitulated endogenous *C15* expression in the antennal disc or leg disc, although the intron 1 enhancer was active in a few cells in the centre of the latter, suggesting it may contain binding sites that contribute to C15 expression in the pre-tarsal region. These results imply that even though *C15* is expressed in these serial homologues, it is regulated by distinct enhancer elements. This modularity may underpin tissue-specific regulatory logic, presumably programmed by different Hox inputs such as Abd-B in the genitalia, and that could have contributed to the diversification of these serially homologous structures from the ‘ground state’ ventral appendage

(6). It would be very interesting to identify the *C15* pre-tarsal and arista enhancers to compare with the genital enhancer that we have identified to understand the differences in *C15* regulation and the GRNs in these developing tissues better. Interestingly, a previous study of bristle number evolution between *D. yakuba* and *D. santomea*, found that the same cis-regulatory change in one enhancer of the *scute* gene affected both sex comb tooth number and hypandrial bristle number in male forelegs and genitalia, respectively (4). Therefore, changes in a single pleiotropic enhancer can also contribute to changes in the morphology of different secondary sexual structures.

It has been shown that the male clasper bristles intercalate with the bristles of the female hypogynium during copulation, and therefore it is likely that the correct pattern of bristles regulated by *C15* is important for mating (5). Furthermore, male terminal structures show great diversity in shape, size and bristle composition, but there are also differences among female structures (6-68). Given the role of *C15* in bristle patterning in male and female terminalia, this gene represents an excellent candidate for differences in the terminal bristle patterns among species.

## Conclusions

We have shown that *C15* is required for the patterning of the bristles in male and female terminal structures. This involves similarities and differences to the regulatory interactions of this transcription factor in other serially homologous appendages, and in particular, the C15-Lim1-Al trio is not used in either male or female terminalia, unlike the antennal aristae and leg pre-tarsal region. Furthermore, *C15* expression appears to be regulated by a different enhancer in the developing male genitalia compared to legs and antennae, which likely unpins the different regulatory logic used in these serial homologues and this modularity may even have facilitated a role for *C15* in the rapid diversification of male genitalia.

## Acknowledgments

This study was funded in part by a BBSRC (BB/X006689/1) grant to A.P.M. and M.D.S.N., a BBSRC DTP studentship (BB/M011224/1) to A.M.R. We thank the Bloomington *Drosophila* Stock Center and Vienna *Drosophila* RNAi Center for fly lines. Microscopy and imaging was carried out in the Centre for Bioimaging at Oxford Brookes University. We thank Gerard Campbell for generously providing antibodies.

## Author contributions

The project was conceived by A.M.R., A.P.M. and M.D.S.N. Experiments were carried out by A.M.R., J.F.J and B.Y.I. Data were analysed by all authors. A.M.R and A.P.M. wrote the paper, assisted by the other authors.

## Supplementary Figures and Tables

**Supplementary Figure 1.**
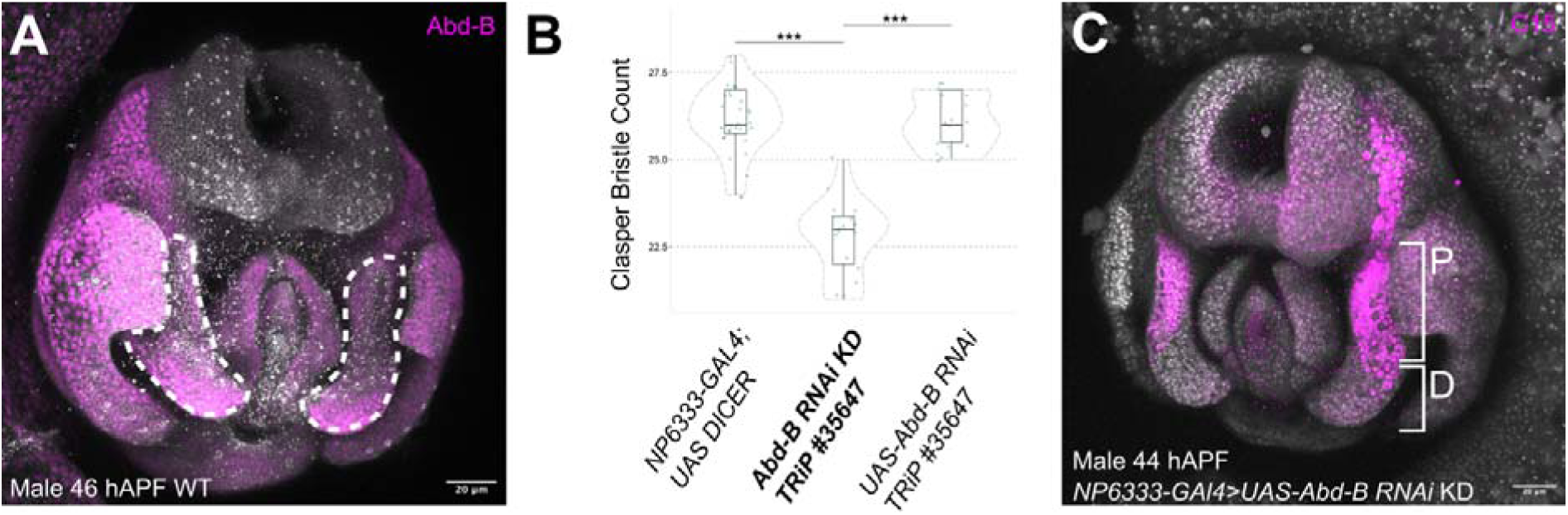
Abd-B expression and regulation of *C15* in the developing male terminalia. (A) Abd-B is expressed throughout the entire male terminalia apart from the cerci. Dashed white lines mark the claspers. (B) RNAi knockdown of *Abd-B* significantly reduced clasper bristle count compared to parental controls. *** p < 0.001 (n > 16). (C) RNAi knockdown of *Abd-B* slightly expanded C15 expression into the distal region of the clasper, as well as at the periphery of the cerci (compare to Figure 2D). P: Proximal, D: Distal clasper regions. All samples were DAPI stained, apart from (A), where anti-DCAD2 staining was used.

**Supplementary Figure 2.**
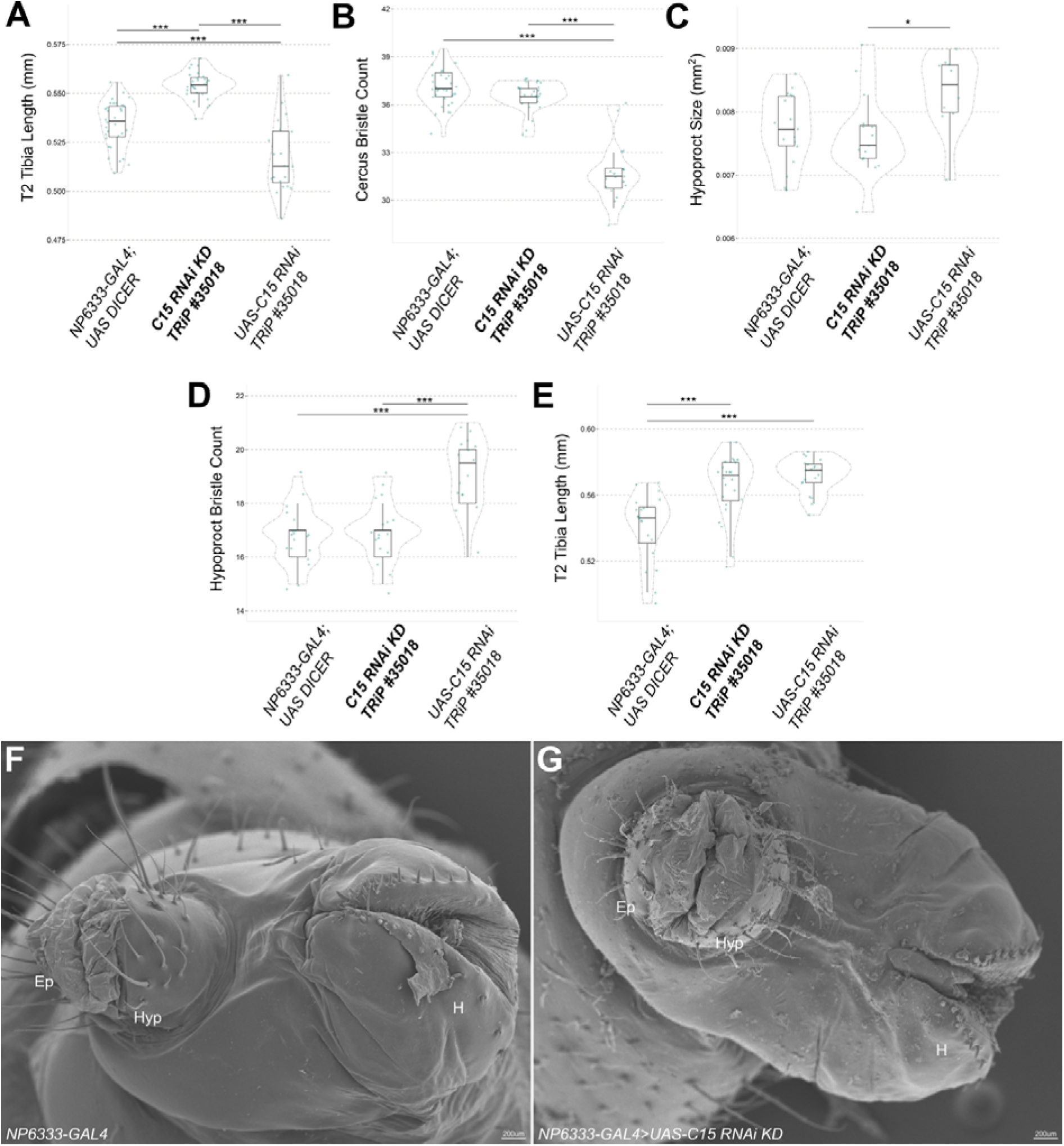
*C15* RNAi knockdown in the developing male and female terminalia and legs. (A) *C15* RNAi knockdown significantly increased male T2 tibia length. (B) *C15* RNAi knockdown did not affect male cercus bristle number. (C) In females, *C15* RNAi knockdown had no significant effect on the hypoproct size (C), hypoproct bristle count (D), or T2 tibia length (E). (F,F’) SEM images of the adult female terminalia in *NP6333-GAL4* parental control (F) and *C15* RNAi knockdown (F’). Epi: Epiproct, Hyp: Hypoproct, H: Hypogynium. *** p < 0.001, and * p < 0.05. n > 9, (Supplementary Tables 1 and 2).

**Supplementary Figure 3.**
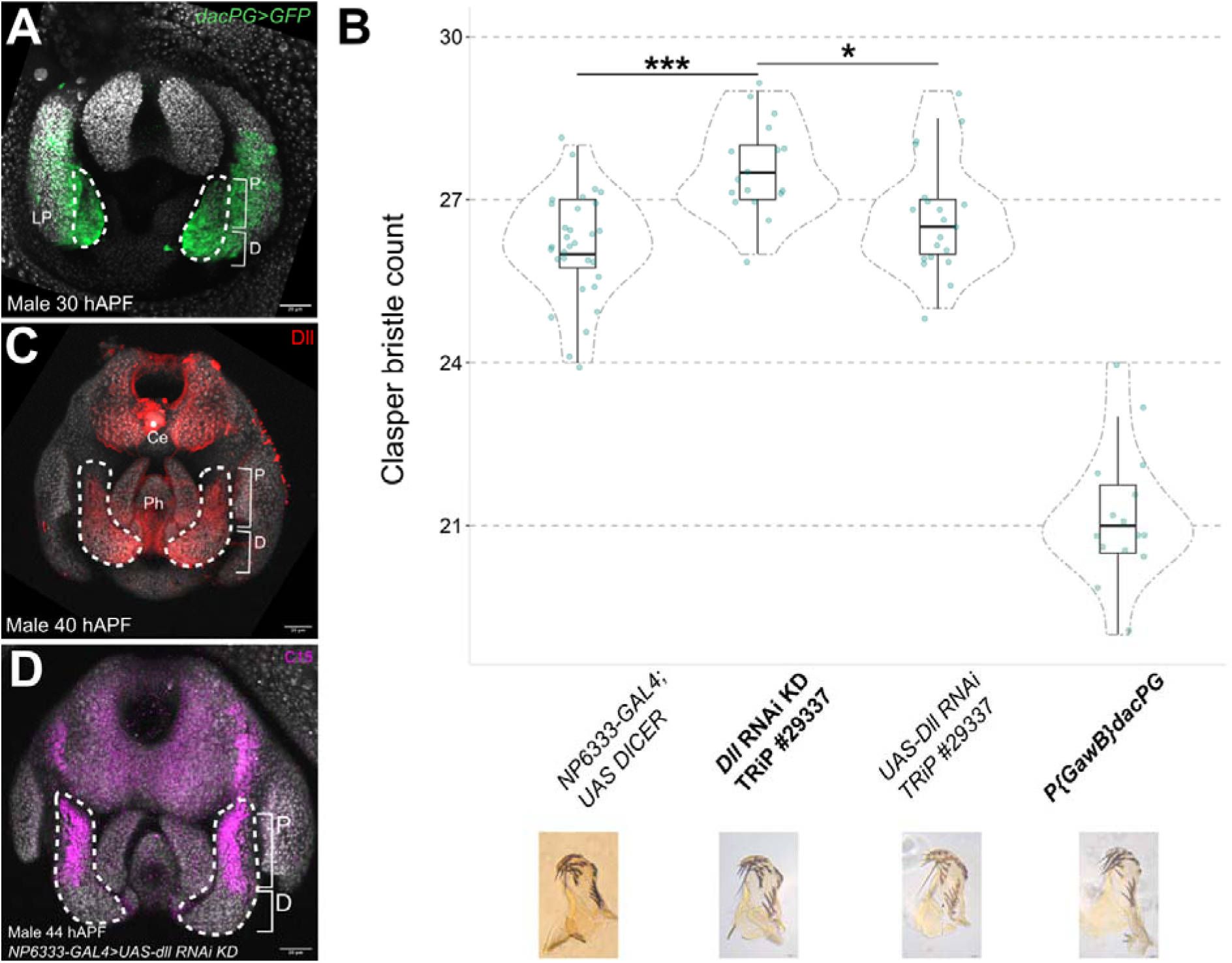
Roles of Dac and Dll during male terminalia development. (A) *dac* is expressed predominantly in the distal clasper regions and in the lateral plate. (B) *Dll* and *dac* are both required for male clasper bristle development. Note the *P{GawB}dacPG* line could not be compared to other genotypes because one parental line was not available for analysis. *** p < 0.001, * p < 0.05 (n > 16 for all lines phenotyped) (Supplementary Table 1). (C) Dll is expressed throughout the claspers, and in part of the phallus, and cerci (1). (D) *Dll* RNAi knockdown did not appear to alter C15 localization. P (proximal) and D (distal) clasper regions. LP: Lateral Plate, Ph: Phallus, Ce: Cercus.

**Supplementary Figure 4.**
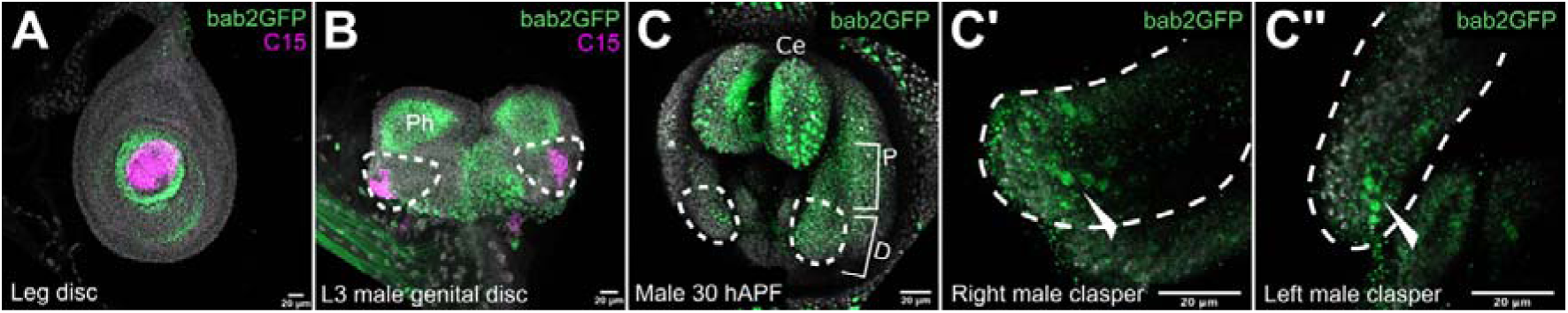
*bab2* expression in developing male terminalia. (A) To verify if the GFP-tagged *bab2* transgenic line recapitulated endogenous *bab2* expression, the leg disc was double stained with anti-GFP and anti-C15. (B) In male genital discs, bab2-GFP localised to the phallus (Ph) and the repressed female primordium. No bab2-GFP expression was observed in the clasper primordia, as marked by C15. (C-C’’) In male pupal terminalia, bab2-GFP was expressed throughout the cerci (Ce), as well as in the distal region of the clasper. All samples were stained with DAPI.

**Supplementary Figure 5.**
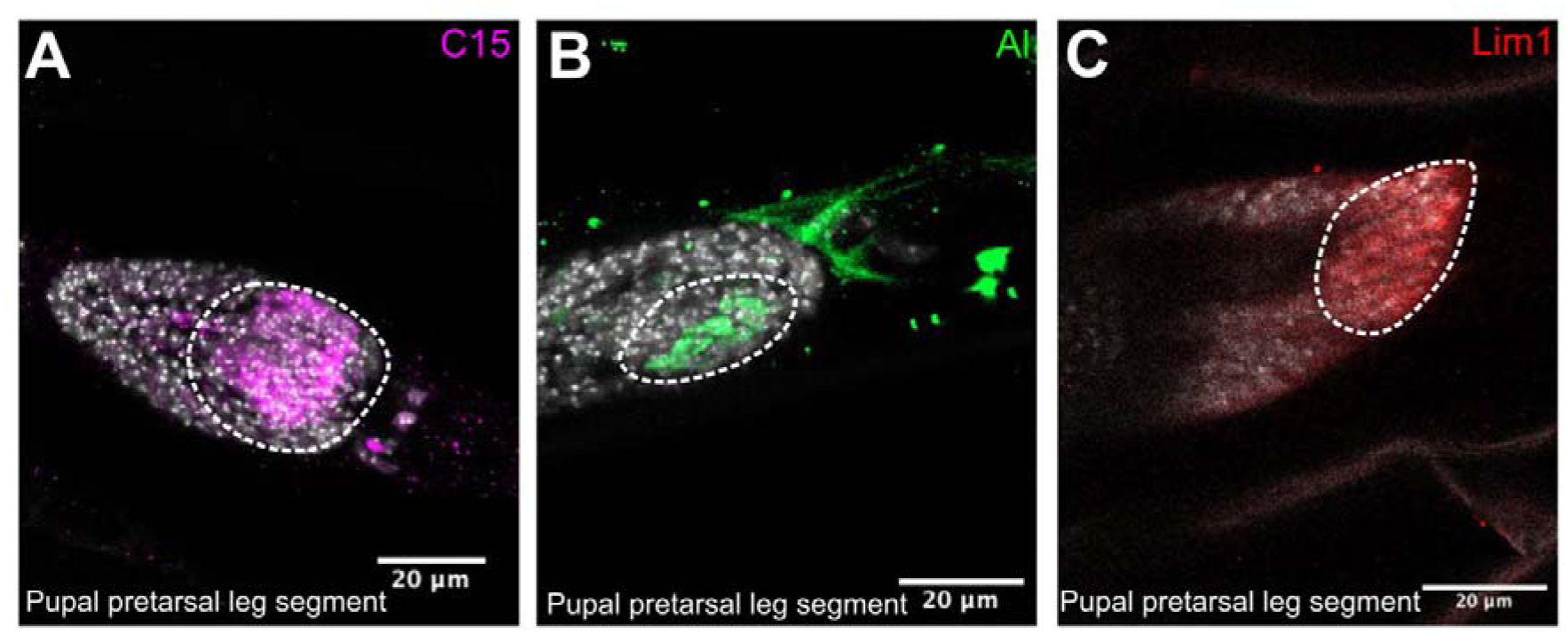
Pupal pre-tarsal leg expression of C15, Al, and Lim1. (A) C15, (B) Al, and (C) Lim1 expression at the pretarsal segment of late pupae legs, where the claw protrudes from. White dashed ovals mark regions of expression. All samples were stained with DAPI.

**Supplementary Figure 6.**
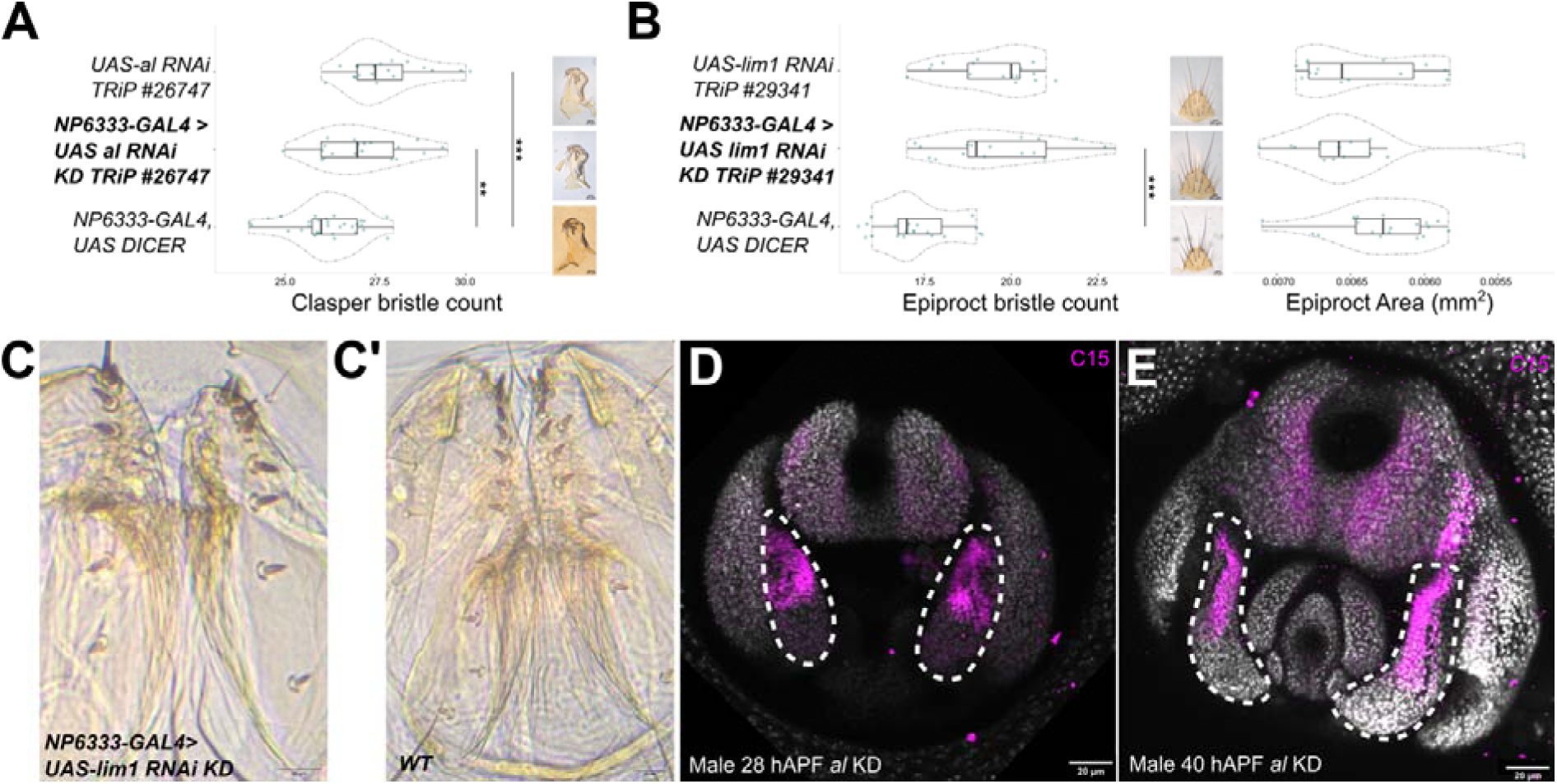
Roles of Al and Lim1 during male and female terminalia development, respectively. (A) RNAi knockdown of *al* did not affect male clasper development. (B) RNAi knockdown of *Lim1* did not affect female epiproct development. (C) *Lim1* RNAi knockdown in the female ovipositor reduced the number of teeth on the hypogynia compared to wild-type females (C’). (D,E) RNAi knockdown of *al* did not affect C15 expression. All samples were DAPI stained. *** p < 0.001, ** p < 0.01 (Supplementary Tables 1 and 2). n > 14 for all lines phenotyped.

**Supplementary Figure 7.**
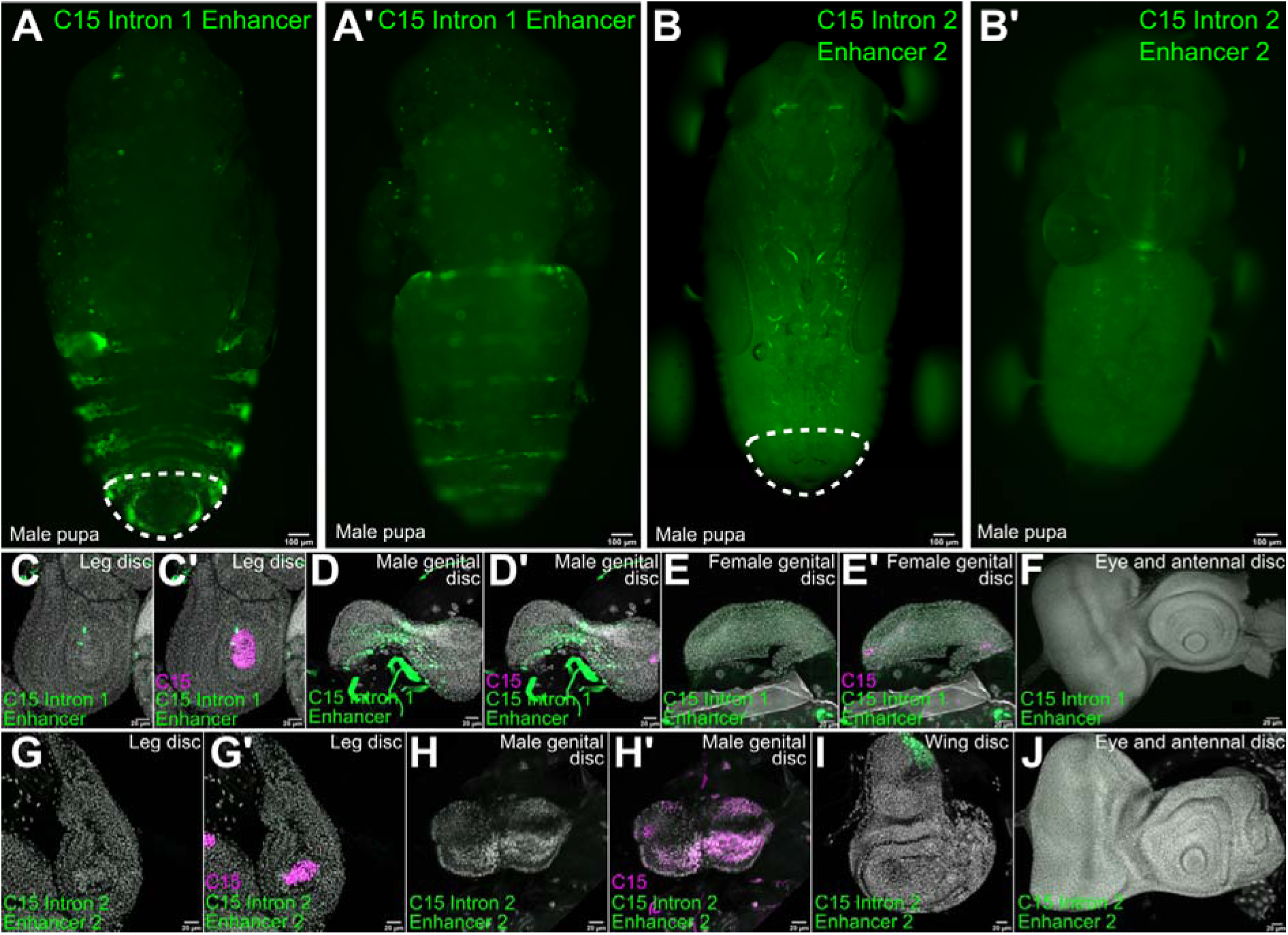
*C15* intron 1 and intron 2 enhancer 2 activity during development. (A,A’) Live imaging of *C15* intron1 enhancer when crossed to *UAS-GFP* in male pupae. *C15* intron 1 enhancer is active in a segmental pattern in the abdomen and in the tissue surrounding the male terminalia. (B,B’) The *C15* intron 2 enhancer 2 is not active in the male terminalia. (C,C’) The *C15* intron 1 enhancer is active in a few cells of the central leg disc and partially overlaps with endogenous C15 expression. (D,D’) This region is also active in the repressed female primordium in the male genital disc but not in the female genital disc (E,E’) nor the eye-antennal disc (F). (G,G’) The *C15* intron 2 enhancer 2 is not active in the leg disc, or the male genital disc (H,H’). (I) This enhancer is active in the notum of the wing disc where endogenous *C15* is expressed (See Figure 7) but not active in the eye-antennal disc (J). All samples were stained with DAPI.

## Supplementary Tables

**Supplementary Table 1. RNAi lines and male phenotypic results.**

**Supplementary Table 2. RNAi lines and female phenotypic results.**

**Supplementary Table 3. Antibodies used in this study.**

**Supplementary Table 4. Transcription factor binding sites predicted by CIS-BP.**

**Supplementary Table 5. Transcription factor binding sites predicted by JASPAR.**

